# Specific expression and function of the Six3 *optix* in serially homologous organs

**DOI:** 10.1101/091793

**Authors:** Amer Al Khatib, Natalia Siomava, Nico Posnien, Fernando Casares

## Abstract

Organ size and pattern results from the integration of two positional information systems. One global, encoded by the Hox genes, links organ type with position along the main body axis. Within specific organs, local information is conveyed by signaling molecules that regulate organ growth and pattern. The mesothoracic (T2) wing and the metathoracic (T3) haltere of *Drosophila* represent a paradigmatic example of this coordination. The Hox gene *Ultrabithorax (Ubx)*, expressed in the developing T3, selects haltere identity by, among other processes, modulating the production and signaling efficiency of Dpp, a BMP2-like molecule that acts as a major regulator of size and pattern. Still, the mechanisms of the Hox-signal integration even in this well-studied system, are incomplete. Here we have investigated this issue by studying the expression and function of the Six3 transcription factor *optix* during the development of the *Drosophila* wing and haltere development. We find that in both organs Dpp defines the expression domain of *optix* through repression, and that the specific position of this domain in wing and haltere seems to reflect the differential signaling profile among these organs. We show that *optix* expression in wing and haltere primordia is conserved beyond *Drosophila* in other higher diptera. Despite the similar expression pattern, *optix* plays different roles in wing and haltere. In the wing, *optix* is required for the growth of the most anterior/proximal region (the “marginal cell”) and for the correct formation of sensory structures along the proximal anterior wing margin. In contrast, in the haltere *optix* is necessary for the suppression of sensory bristles without any noticeable effect on organ growth. Therefore, *optix* shows an organ-specific function. Beyond dipterans, *optix* expression in the anterior wing has been shown also in butterflies. We propose that the ancestral role of *optix* might have been structural in the anterior wing. Once expressed in the wing, *optix* expression had been re-deployed for wing spot formation in other parts of the wing of Heliconius butterflies.

## INTRODUCTION

During development, several positioning systems inform cells of their location. First, the Hox code defines position along the anterior-posterior axis. In insects, this system generates segmental diversity along this body axis. Next, in each segment, cells within an organ primordium obtain positional information from local signaling centers (Mann and Carroll, 2002). The *Drosophila* wing and haltere primordia constitute a paradigm where the integration of these two positional systems has been intensively investigated. In the *Drosophila* thorax (T), the second and third segments develop two serially homologous organs: the wing (in T2) and the haltere (in T3). The specific expression of the Hox gene *Ultrabithorax (Ubx)* in T3 is responsible for the specific morphology of the haltere, a small, modified wing that acts as balancing organ during *Drosophila* flight. Mutants that cause the loss of Ubx during haltere development cause its transformation into wing (Lewis, 1978), while ectopic expression of Ubx in the developing wing results in wings transformed into haltere-like appendages (Gonzalez-Gaitan et al., 1990).

One of the major organ-positioning systems in the wing and haltere primordia -or “imaginal discs” – is a stripe of cells that bisects the disc and produces a BMP2 ligand, encoded by *decapentaplegic* (*dpp*). From this stripe, Dpp generates a signaling gradient that patterns the anterior-posterior axis of the appendage (Restrepo et al., 2014). In the wing, which is the ground state of the dorsal appendage, the Dpp gradient activates the nested expression of a number of target genes at different concentrations, such as *spalt (sal*, referring collectively to two highly related paralogous genes, *sal-m* and *sal-r)* and *optomotor blind (omb)* (Nellen et al., 1996). *sal* and *omb* activation is indirect, though, by Dpp signaling repressing *brinker (brk)*, itself a repressor of the Dpp pathway (Campbell and Tomlinson, 1999; Winter and Campbell, 2004). In this way, Dpp controls the positioning of central pattern elements, such as the wing veins (de Celis et al., 1996; Sturtevant et al., 1997). In addition to patterning, Dpp signaling controls organ growth, so that mutants that lack Dpp signaling result in very reduced winglets (Posakony et al., 1990).

In the haltere, *Ubx* modifies the wing developmental program in two ways. First, as a transcription factor, Ubx regulates the expression of some targets. For example, Ubx represses *sal* expression (Weatherbee et al., 1998). Second, *Ubx* modifies the shape of the Dpp-generated signaling gradient indirectly, by controlling the expression of proteoglycans required for Dpp dispersion (Crickmore and Mann, 2006; de Navas et al., 2006b). Globally, these modifications of Dpp signaling and target gene activation by Ubx have been related to the size and patterning differences between halteres and wings.

Although Dpp signaling generates a signaling gradient that spans the whole wing pouch and its activity is required throughout the wing, so far only centrally expressed target transcription factors have been described. Here we report the functional involvement of the transcription factor Optix/Six3 in patterning of the most anterior region of the wing and the haltere. In both discs, *optix* expression is anteriorly restricted by Dpp signaling, although in the wing the precise expression boundary may be set with the collaboration of wing specific Dpp targets, such as *sal.* We show that *optix* shows organ-specific functions: in the wing, it is necessary for the growth of the anterior/proximal wing (“marginal cell”) and the development of wing margin sensory bristles, while in the haltere *optix* is required for the suppression of sensory bristle formation. Overexpression of *optix* in the entire wing pouch affects only anterior wing development, suggesting that other parts of the wing cannot integrate ectopic Optix input. This observation may provide a mechanistic explanation for a widespread redeployment of *optix* expression in wing spot formation in various butterfly species.

## MATERIALS AND METHODS

### Fly strains and genetic manipulations

Two *optix:GFP* lines were examined (Sarov et al., 2016) (318456/88 and 318371/10042). Expression of both lines was qualitatively similar, but the signal of *optix:GFP 318371/10042* was strongest and was used for all further studies (and referred to as “*optix*.GFP”).

Reporter strains used were: *hh^P30^-lacZ* (Lee et al., 1992) and *ap-lacZ* (Cohen et al., 1992. The TM2 balancer carries the *Ubx*^*130*^ allele (Flybase: http://flybase.org), which expresses a partial transformation of the haltere into wing, including the presence of small triple row-like bristles.

The UAS/GAL4 system (Brand and Perrimon, 1993) was used for most gain- and loss-of-function assays. Given that we focused our investigation in the pouch region, we used as wing specific driver *nubbin-GAL4 (nub-GAL4)* (Calleja et al., 1996). As reporter of the expression of *optomotor-blind (omb)*, the *omb-GAL4* strain (3045, Bloomington Stock Center) was crossed to *UAS-Cherry-RFP* (27391, Bloomington Stock Center). *UAS-optixRNAi* (33190, Bloomington Stock Center) and *UAS-puntRNAi* (37279, Vienna Drosophila Resource Center) were used for gene-specific knockdown induction, and UAS-OptixS1 (26806, Bloomington Stock Center) for *optix* ectopic expression experiments.

To generate *tkv* loss-of-function clones through mitotic recombination (Xu and Rubin, 1993), we crossed 1096-GAL4,UASflp;FRT40arm-/acZ/Cy0 females to *tkv*^*a12*^ FRT40A/CyO males. *tkv*^*a1*^ is a tkv-null allele. In this experiment, the *bxMS1096-GAL4* line (“1096-GAL4" (Milan et al., 1998) drives UAS-flipase throughout the wing disc to induce mitotic recombination clones in this organ specifically. *optix:GFP* was introduced in these genotypes by standard genetic techniques. All crosses were raised at 25°C, except in the case of UAS-RNAi experiments, which were transferred to 29°C 48 hours post-fertilization (hpf) to maximize the penetrance of the knockdowns.

The Mediterranean fly strain *Ceratitis capitata* Egypt II was obtained from the FAO/IAEA Agriculture and Biotechnology Laboratory (Seibersdorf, Vienna, Austria) and reared at 28°C and 55 ± 5% RH. The house fly strain *Musca domestica* ITA1 was collected in Italy, Altavilla Silentia in 2013 (Y. Wu and L. Beukeboom, GELIFES, The Netherlands) and kept at room temperature (22±2°C) on wheat bran-based food.

### *in situ* hybridization

Images of *optix* mRNA expression in wing and haltere discs are unpublished data kindly shared by P. Tomancak (MPI-CBG, Dresden) and C. Dahmann (Technische Universität, Dresden), obtained using probes and methods described in (Tomancak et al., 2007).

Orthologous *optix* sequences for *C. capitata* and *M. domestica* were obtained by NCBI BLAST starting with the *Drosophila optix* sequence. Fragments were amplified with gene specific primers for *C. capitata*: forward: GACCGACGGAGGGCAAACATCCTCC and reverse: GTTCAAGCTATGCGCCTGTGCCGGC; and for *M. domestica:* forward: GACCGACGGAGGGTAAACAACCTCAAC and reverse: CGGCCGCATCCAGTTTAAACGAAGGC. The digoxigenin (DIG)-labeled antisense RNA probes were synthesized from purified PCR products by using the DIG RNA Labeling Mix, T7-RNA Polymerases, and Protector RNase Inhibitor (Roche Applied Science, Mannheim, Germany) and fragmented to an average length approx. 200 bp by adding an equal amount of sodium carbonate buffer (80 mM NaHC03, 120 mM Na2C03, pH 10.2) followed by an incubation at 60°C. Fragmented probes were diluted with HybeA buffer (50% formamide, 0.1 μg/μl sonicated salmon sperm DNA, 50 μg/ml Heparin, 5 x SSC and 0.1% Triton X-100, in PBS) and used for *in situ* hybridization. Wing and haltere imaginal discs were dissected from *C. capitata* and *M. domestica* third instars and fixed in 4% PFA for 30 min. After fixation, samples were washed three times with PBT for 20 min, rinsed once with 1:1 HybeA:PBT, and quickly washed three times with HybeA. Pre-hybridization was performed in HybeA at 65°C for 1 h. Preheated and chilled down probes were added to samples and hybridized overnight at 65°C. On the next day, probes were discarded, samples were washed three times with preheated HybeA at 65°C for 20 min and one time with 1:1 HybeA:PBT, and incubated with 1ml of anti-DIG-AP antibody (Anti-Digoxigenin-AP, Fab fragments, Roche Applied Science, Mannheim, Germany, diluted 1:2000 in PBT) at room temperature for 1 h. Antibodies were removed, samples were washed three times with PBT for 20 min and three times with a freshly prepared detection NBT buffer (100 mM Tris-HCl with pH 9.5, 100 mM NaCl, 50 mM MgCl2, 0.1% TritonX-100, in water) for 5 min. After the last washing step, samples were transferred into glass wells and the detection buffer was replaced with the staining solution (1 ml NBT buffer, 4.5μl NBT (Nitrotetrazolium Blue chloride, Carl Roth GmbH & Co KG, Karlsruhe, Germany, 50 mg/ml in 70% DMF), and 3.5μl BCIP^®^ (5-Bromo-4-chloro-3-indolyl phosphate disodium salt, SIGMA-ALDRICH^®^ Chemie GmbH, Munich, Germany, 50 mg/ml in 100% DMF)). Samples were incubated in the dark at room temperature. The staining reaction was stopped by washing samples three times with PBT for 10 min each.

### Immunofluorescence and confocal imaging

Immunofluorescence in wing and haltere imaginal discs was carried out according to standard protocols. Primary antibodies used were: mouse anti-GFP (Molecular Probes), rabbit anti-GFP (Molecular Probes), rabbit anti-pSmad3 (AbCam), anti-Sal (gift of C. Sánchez Higueras and J. Hombría, CABD, Seville) and anti-Ubx (#FP3.38, Iowa University Hybridoma Bank). *lacZ* reporters were detected using a rabbit anti-β-galactosidase antibody (#55976, Cappel). Appropriate Alexa Fluor-conjugated secondary antibodies were used. For experiments that were used for fluorescence intensity quantification, confocal settings were kept constant, so that fluorescence intensity could be compared across discs. After immunostaining, samples were imaged using a Leica SPE confocal microscope. Images were processed with ImageJ (NIH).

### Quantification of gene expression profiles

Expression profiles for *optix:GFP*, PMad (monitored using a cross-reacting antibody against P-Smad3), *sal* and *omb* (monitored in *omb>RFP* larvae) were obtained using the “Plot profile” function of ImageJ (NIH). Within each experiment, confocal settings were maintained constant so intensity profiles would be comparable. Intensity profiles were expressed in arbitrary units.

## RESULTS

### The Six3 gene *optix* is differentially expressed in the wing and haltere discs

*Optix* transcription, detected using RNA *in situ* hybridization, is found in both the wing and haltere imaginal discs of late third instar (L3) larvae (Organista et al., 2015; Seimiya and Gehring, 2000) (Figure 1A) in the pouch regions of both discs. These pouch regions give rise to the wing proper and the distal haltere’s article (capitellum), respectively (Figure 1C; (Cohen, 1993). To examine the expression of *optix* in detail, we used an Optix:GFP line (Sarov et al., 2016) that recapitulates *optix* expression (Figure 1B). We first mapped optix-expressing domains in the wing and haltere discs relative to the anterior-posterior (AP) and dorso-ventral (DV) boundaries. We used *apterous (ap; ap-Z)*, as a D marker, and *hedgehog (hh; hh-Z)* as a P marker. Relative to the DV axis, *optix* straddles symmetrically the DV boundary in both wing and haltere discs (Figure 1D,E). *optix* expression is restricted to the A compartment in both discs (Figure 1F,G). However, the position that the *optix* domain occupies along the AP axis is different: in the wing, *optix* is restricted to the anterior-most region of the pouch, while in the haltere it occupies a more central position, closer to the AP border.

**Figure 1.**
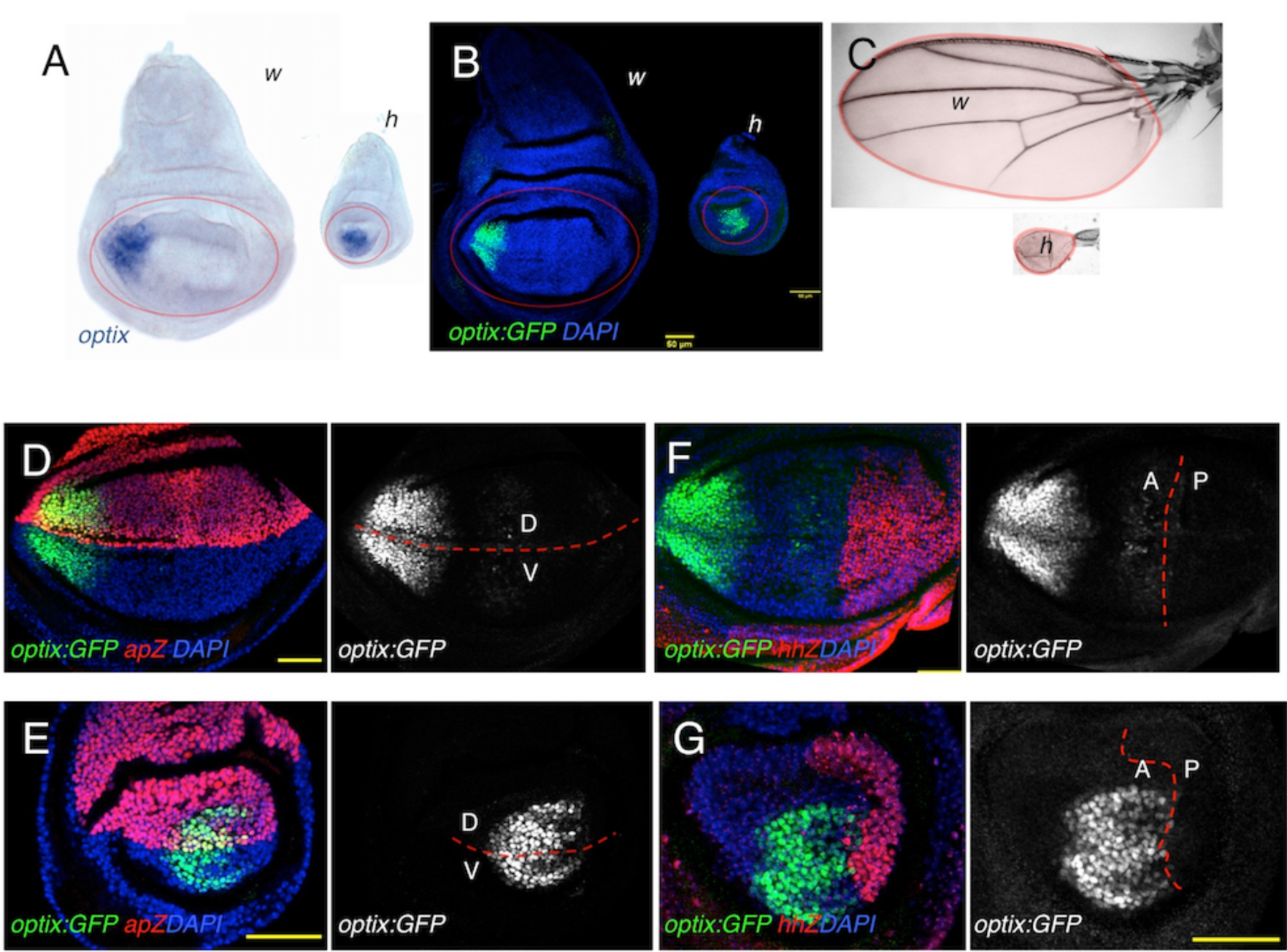
*optix* expression in wing and haltere discs relative to positional references. (A,B) *optix* expression detected by in situ hybridization (A) or monitored by the Optix:GFP strain (B) in late third instar wing (w) and haltere (h) imaginal discs. The pouch region in wing and haltere discs (outlined in A) gives rise to the wing blade and haltere capitellum, respectively, colored in red in (C). (D,G) *optix:GFP* expression relative to *ap-Z* (D,E) and *hh-Z* (F,G) in wing (D,F) and haltere (E,G) discs. Bar: 50μm.

### *optix* expression in wing and haltere discs is conserved within higher diptera

*Drosophila melanogaster* is a highly derived dipteran. To test whether the wing and haltere expression of *optix* is conserved beyond *Drosophila*, we analyzed the expression pattern of the *optix* homologues in two Schizophoran fly species: *Ceratitis capitata* (Tephriditae) and *Musca domestica* (Muscidae). Using *in situ* hybridization, *optix* is detected in equivalent patterns in wing and haltere discs of these two species (Figure 2A-D), indicating that *optix* expression pattern is conserved during wing and haltere development within higher diptera.

**Figure 2.**
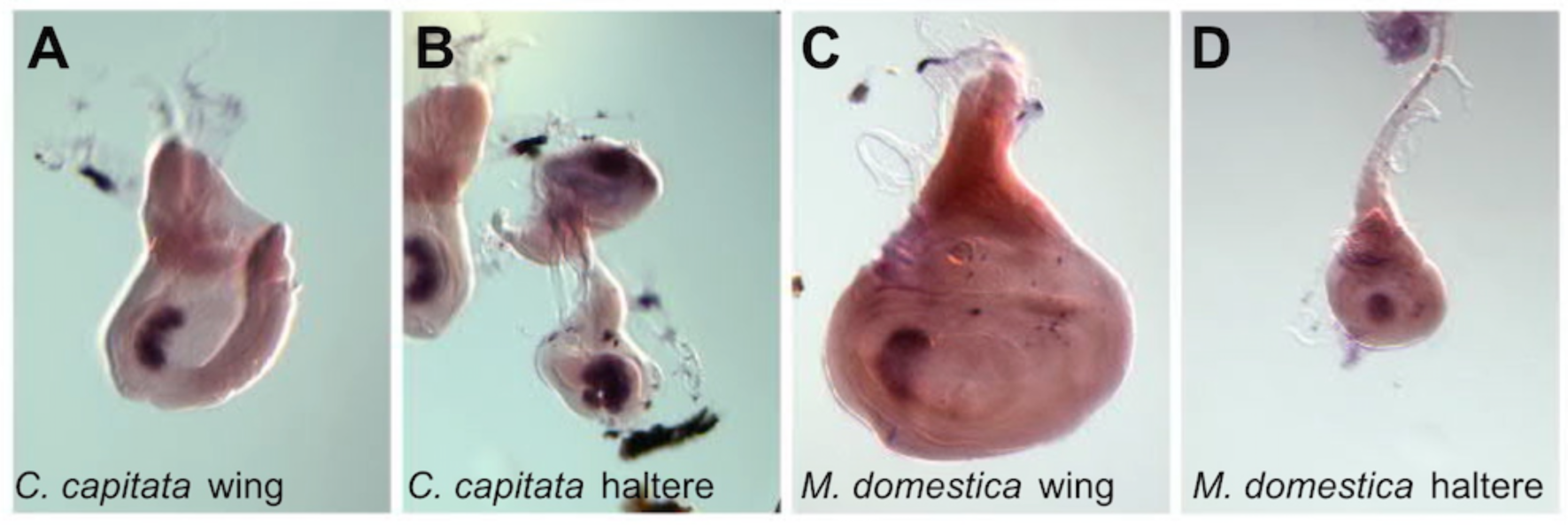
*optix* expression in other dipteran species. *in situ* hybridization detection of *optix* in wing and haltere discs of third instar larvae from *C. capitata* and *M. domestica.* In both species, the expression pattern in wings and haltere discs is very similar to the one seen in *D. melanogaster.*

### *optix* plays different roles in wing and haltere development

In order to determine the role played by *optix*, we drove an optix-RNAi to the distal wing and haltere discs, using the *nubbin-GAL4 (nub>)* driver (Figure 3A). The adult wings of *nub-GAL4; UAS-optixRNAi (“nub>optixRNAi”)* flies were bent and smaller than those of their *nub>+* siblings (Figure 3B). This phenotype seemed to be mostly due to a much shorter longitudinal vein 2 (L2) and a reduction of the wing blade area anterior to this vein (the so-called “marginal cell”, in between veins 1 and 2; Figure 3C,D) to about ¼ the normal area. The density of trichomes in the wing tissue, that can be used as a proxy for cell size, is very similar in the marginal cell of *nub>+* and *nub>optixRNAi* (density of trichomes in *nub>optixRNAi* is 0.93 times that of *nub>+* controls). Therefore, the area reduction of the marginal cell is the result of reduced growth. This area reduction is accompanied by a loss of margin sensory bristles (Figure S1). In contrast, in the haltere *optix* attenuation did not result in any noticeable size difference. Instead, *nub>optixRNAi* halteres developed extra bristles in the capitellum, similar to those found in *Ubx+/-* heterozygous individuals (Figure 3E-G). However, while the bristles in *Ubx+/-* appeared along the anterior region of the haltere (which in this genotype is larger than wild type), in *nub>optixRNAi* the bristles develop in a more medial position on the ventral side of the haltere (Figure 2E-G).

**Figure 3.**
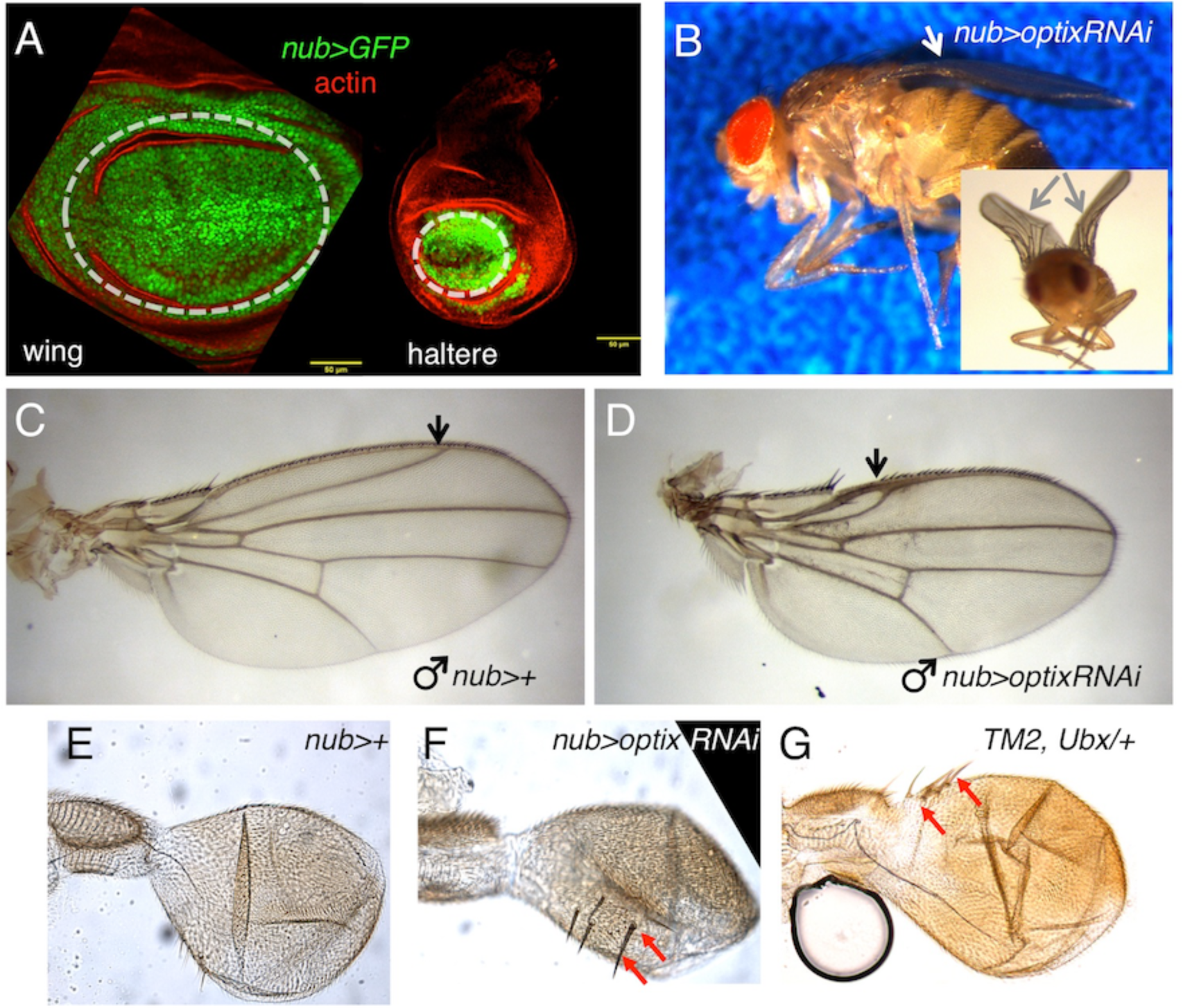
*optix* knock-down affects wing and haltere development. Expression of *optix-RNAi* was driven by the *nubbin-GAL4 (“nub>”*) driver. In wing and haltere discs (A), *nub>* drives expression in the central portion of the discs, including the wing and haltere pouches (outlined by the dashed white line), as observed by GFP expression in *nub>GFP* discs. Discs were counterstained with rhodamine-phalloidin, which marks filamentous actin (“actin”). Adult *nub>optixRNAi* male. The inset shows a frontal view. Wings show a characteristic bent (arrows). Wings from control *(nub>+;* C) and *nub>optixRNAi* (D) males. L2 (arrow) is shorter and the wing area in between the margin and L2 is severely reduced. Halteres from adult *nub>+* (E), *nub>optixRNAi* (F) and *TM2, Ubx/+* (G) males. Red arrows point to ectopic bristles.

So far, our results indicated that *optix* is expressed at a different position along the AP axis of wing and haltere primordia, where it plays organ-specific functions. We decided to investigate next the mechanism responsible for the precise AP positioning of the *optix* domain.

### *optix* expression is complementary to high Dpp signaling levels

In the wing and haltere primordia AP information is conveyed by Dpp (Decapentaplegic), a BMP2-like molecule. Dpp is produced at a stripe along the AP axis from where it diffuses, creating a signaling gradient (Restrepo et al., 2014). Cells regulate gene expression according to the signaling levels they perceive –i.e. according to their position. The read-out of this Dpp signaling gradient is the phosphorylated form of the transcription factor Mad (pMad) (Sekelsky et al., 1995). However, the shape of the Dpp signaling gradient (i.e the pMad profile) differs between wing and haltere discs. This difference has been shown to be the result of Ubx regulating the production and spread of Dpp in the haltere (Crickmore and Mann, 2006; de Navas et al., 2006a). In addition, Ubx regulates directly the output of the Dpp pathway, for example, by repressing Dpp’s target *spalt (sal)* in the haltere (Barrio et al., 1999; Galant and Carroll, 2002; Weatherbee et al., 1998). Since we have found that *optix* was expressed more laterally in the wing than in the haltere pouch, we asked whether the position of *optix* relative to the Dpp signaling gradient was also different. To do this, we stained *optix:GFP* discs for pMad, using a crossreacting antibody against the mammalian pSMAD3. We confirmed previous observations indicating that that the maximum intensity of the pMad gradient was lower in the haltere, indicating a weaker Dpp signaling in the haltere compared to the wing (Figure 4; (Crickmore and Mann, 2006). Within this gradient, *optix* was excluded from regions of high and intermediate signal (100-600 arbitrary units) in the wing (Figure 4B) as well as in the haltere, where *optix* was displaced relatively to the pMad gradient, so that its expression was excluded from the peak of pMad signal (100-150 arbitrary units) (Figure 4A-D). This meant that the expression domain of *optix* relative to the Dpp signaling intensity was similar in both discs despite their *optix* domains being located far from (in the wing) or adjacent to (haltere) the AP border (Figure 1). The complementarity of expression suggested that the positioning of *optix* expression was set by Dpp repressing *optix.*

**Figure 4.**
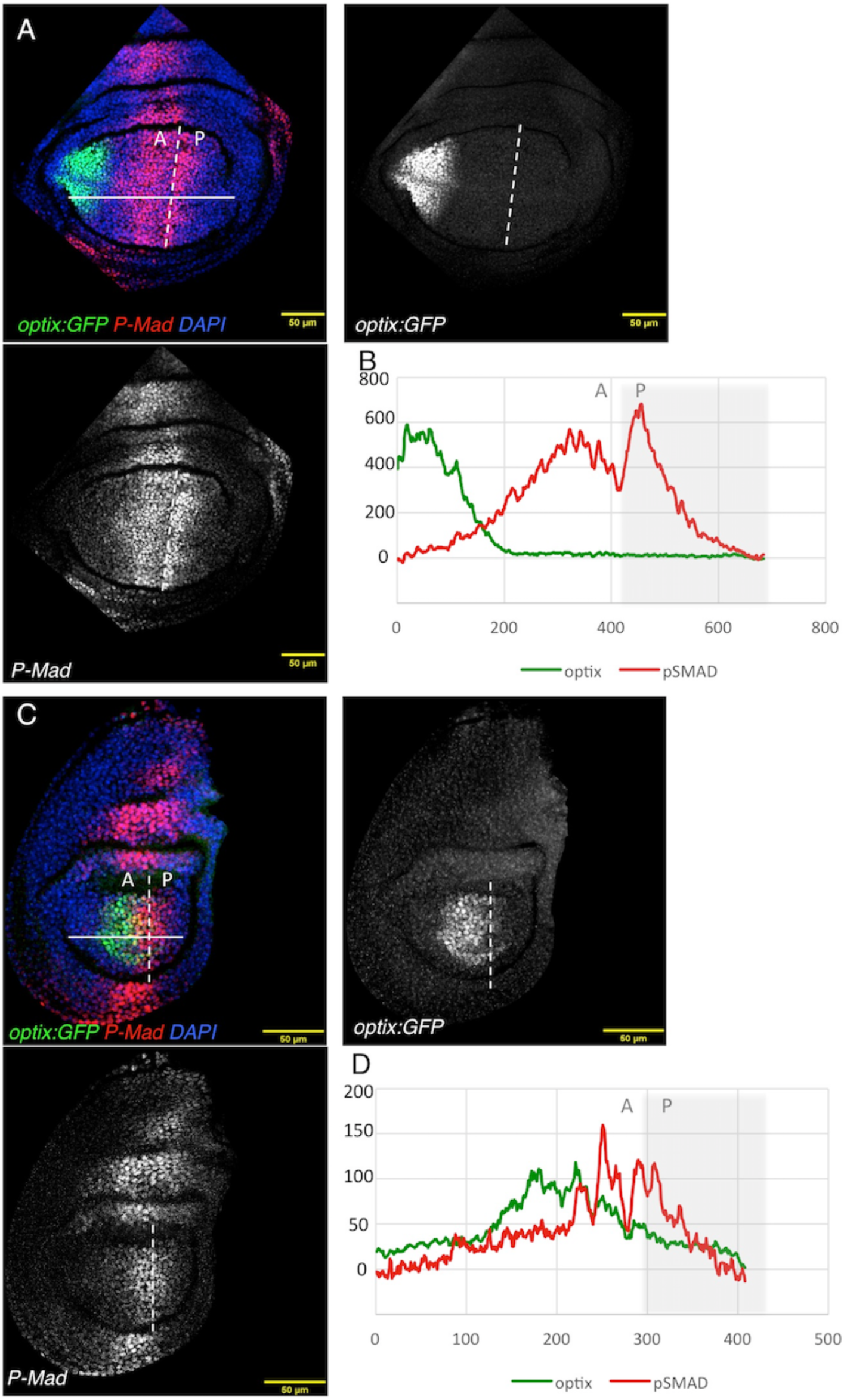
*optix* expression relative to the pMad signaling profile. *Optix:GFP* wing (A) and haltere (C) discs stained for pMad (red), and counterstained with the nuclear marker DAPI (blue). *>GFP*> and pMad profiles for wing (B) and haltere (D) discs, across the white lines, are shown. Fluorescence intensity in arbitrary units. “A” and “P”: “anterior” and “posterior” compartments, respectively. The AP border is marked by the white dashed line.

### Dpp signaling represses *optix* and sets the limits of its expression domain

Dpp signaling could be repressing *optix* directly or indirectly, through some of its targets. To test the hypothesis of a repressor role for the Dpp pathway, we first attenuated the expression of the Dpp type II receptor *punt* (Letsou et al., 1995; Ruberte et al., 1995) using a *punt*-specific RNAi (Figure 5). In *nub>puntRNAi* wing and haltere discs the *optix* domain extended towards the disc center. In these discs, the pMad signal from the pouch is absent, confirming the blockade of the signaling. This was especially noticeable in the wing disc (Figure 5A,B). Although this result suggested a repressive action of the Dpp pathway on *optix*, it could not rule out that low levels of Dpp signaling could be activating *optix* as, in our experiment, *punt* levels had been attenuated using an RNAi. To unambiguously assess the role of Dpp in the regulation of *optix*, we induced loss-of-function clones of *tkv*, the Dpp type I receptor (Brummel et al., 1994; Nellen et al., 1994). In these clones, that grow poorly and tend to extrude from the epithelium (Burke and Basler, 1996), we detected depression of *optix:GFP* in clones all along the AP axis of the wing (Figure 5C). As these clones cannot transduce the Dpp signal, we conclude that Dpp signaling is a repressor of *optix.*

**Figure 5.**
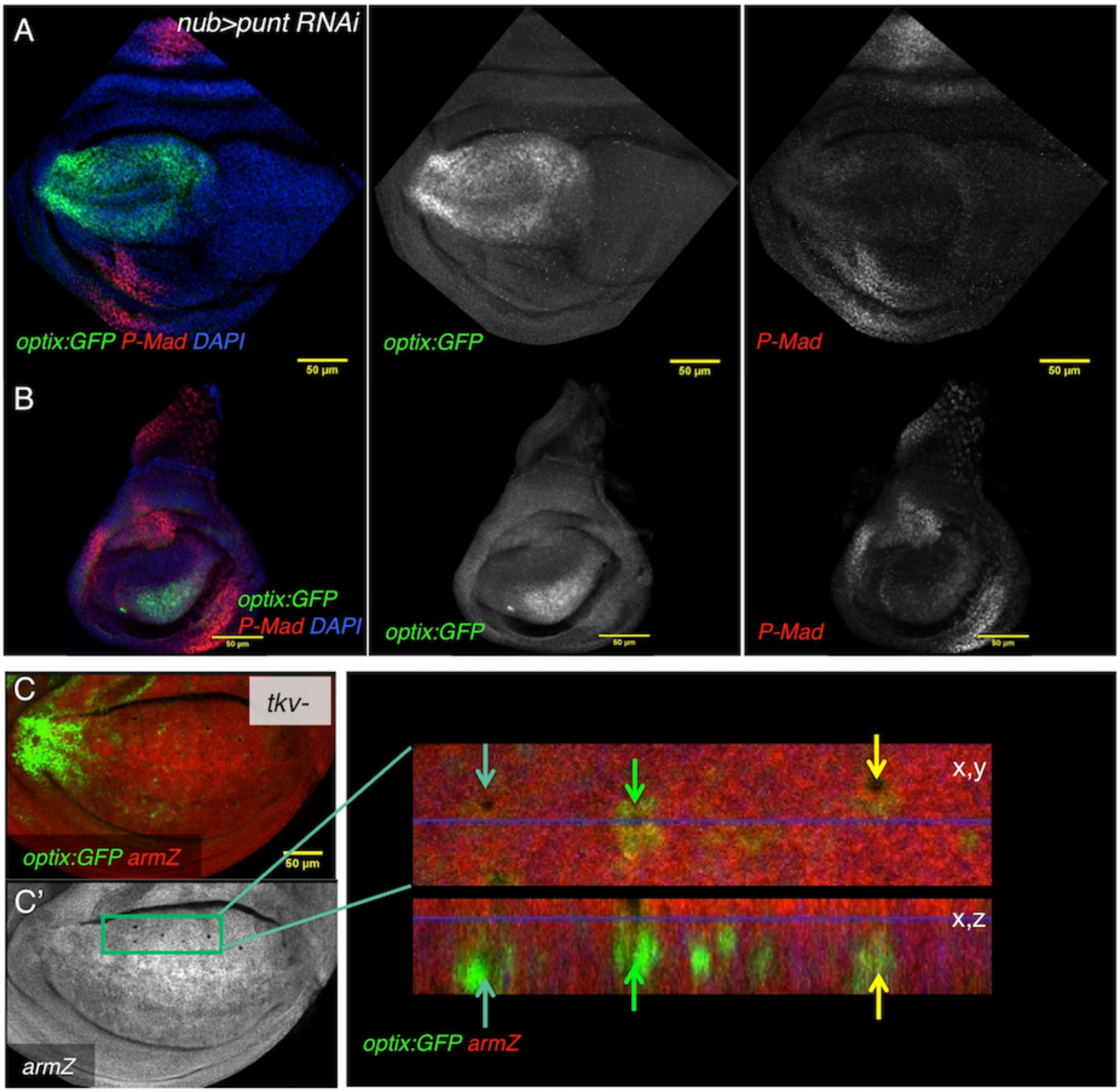
Dpp signaling represses *optix:GFP*. *nub>optixRNAi; optix:GFP* wing (A) and haltere (B) discs. *optix:GFP* signal expands. In these discs, the pMad signal characteristic of the pouch is lost. Discs are counterstained with the nuclear marker DAPI. (C) Wing disc containing tkv-mutant clones. The clones are marked by the absence of *armZ* (C’ shows the *armZ* signal alone). As these clones tend to sort out from the epithelium, they are detected as small “holes” in an apical view. To the right, a detail of the same wing pouch, showing an apical view (“x,y”) and a confocal z-optical section (“x,z”). Arrows mark clones in the (x,y) and (x,z) planes. All these clones, which are small and sort out basally, express *optix:GFP.*

To examine the possibility that the repressive action was exerted through some of its targets, we analyzed the expression of *optix* in wing and haltere discs relative to two known Dpp target transcription factors, *spalt (sal)* and *optomotorblind (omb)* (de Celis et al., 1996; Grimm and Pflugfelder, 1996; Kim et al., 1996; Nellen et al., 1996; Sturtevant et al., 1997). In the wing, the *sal* and *optix* domains are separated by an intermediate zone and do not overlap (Figure 6A,A’). In the haltere pouch, though, *sal* is not expressed (Figure 6B,B’) and yet, as we showed above, *optix* expression is excluded from the regions of intermediate/high Dpp signal. These results do not rule out *sal* as an *optix* repressor in the wing, but suggest that it cannot be the sole repressor, as it is absent from the haltere. Indeed, Organista et al. (2015) have shown that in *sal-*mutant wing discs, *optix* expression extends towards the disc’s center, but does not reach the AP boundary, indicating that additional Dpp-dependent mechanisms for *optix* repression must exist. Next, we analyzed the expression of *optix* relative to *omb* in *omb-GAL4; UAS-cherry-RFP.* While in the wing *optix* and *omb* were complementary to one another (Figure 6C,C’), in the haltere we detected significant overlap between both genes (Figure 6D,D’). Therefore, *omb* does not seem to fulfill the repressor role either, because of its coexpression with *optix* in the haltere.

**Figure 6.**
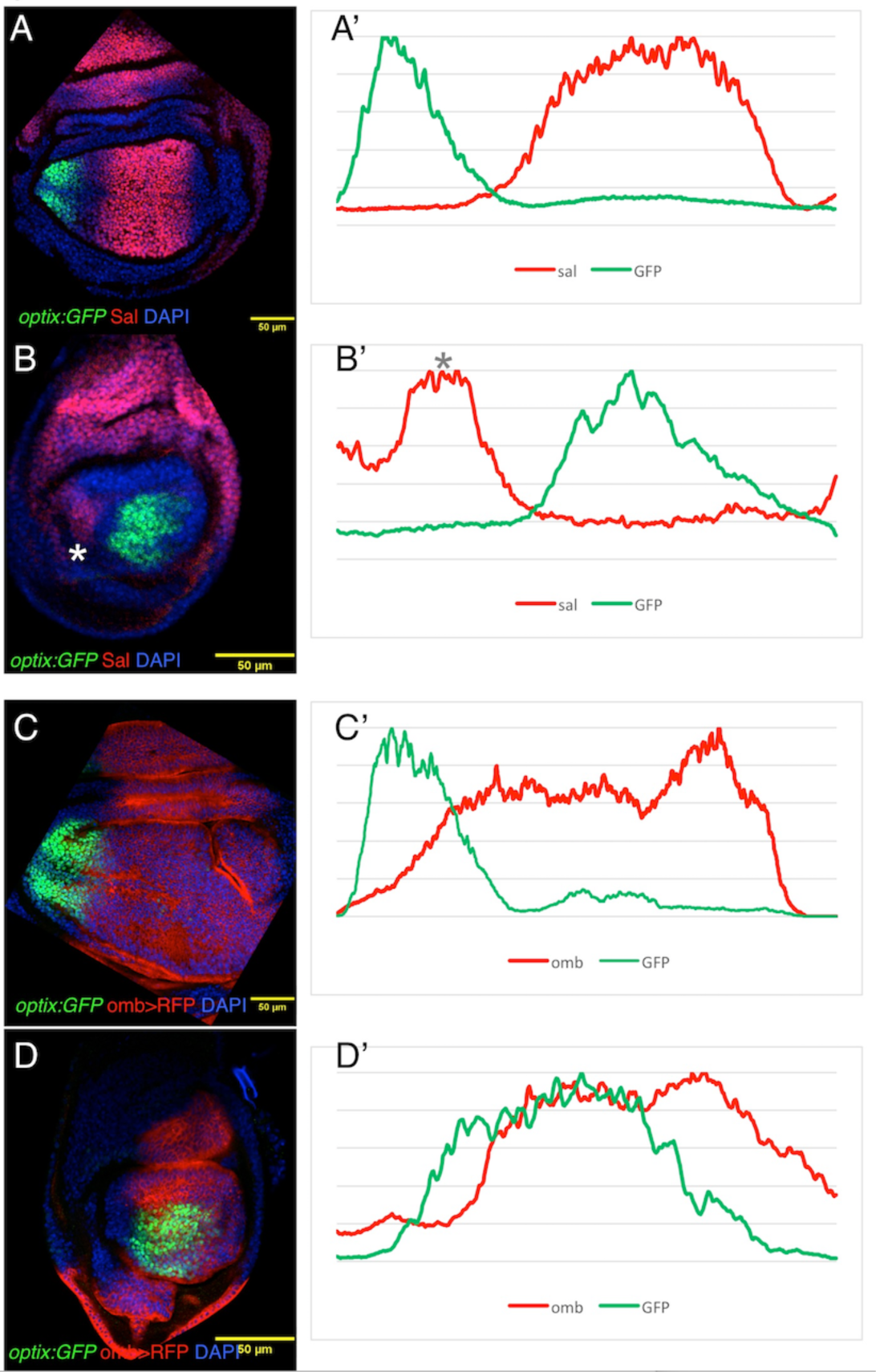
Spatial relationship between *optix:GFP* and the Dpp targets *sal* and *omb.* (A,B) wing (A) and haltere (B) discs of the *optix:GFP* line stained for Sal, and counterstained with the nuclear marker DAPI. (A’) and (B’) are expression profiles through the pouch regions of the same wing (A’) and haltere (B’) discs. (C,D) wing (C) and haltere (D) discs from *omb>RFP;;optix:GFP* larvae, and counterstained with the nuclear marker DAPI. (C’) and (D’) are expression profiles through the pouch regions of the same wing (C’) and haltere (D’) discs. The asterisk in (B) and (B’) marks a very proximal domain of *sal* expression in the haltere.

### *optix* seems to function parallel or downstream of *Ubx*

We find that there is a two-fold relationship between *optix* and *Ubx.* On the one hand, *Ubx* is responsible for the modification of Dpp positional system in the haltere, which then sets *optix* domains along the AP position. On the other, *optix* expression is required in the haltere to suppress the formation of sensory bristles in this organ –a function known to be exerted by Ubx (Garcia-Bellido and Lewis, 1976; Weatherbee et al., 1998). In principle, this latter phenotype could be produced if *optix* were required for either *Ubx* expression or function. Alternatively, *optix* could be required for one of Ubx’s activities: the suppression of bristle development. We tested the first possibility by examining *Ubx* expression in *nub>optixRNAi* haltere discs, stained with an anti-Ubx antibody. In these discs we did not observe any change in Ubx protein levels relative to controls (Figure S2A,B). This result was not unexpected, as a reduction in *Ubx* levels would have led, in addition to extra bristles, to an increase in haltere size, something we do not observe in *nub>optix RNAi* individuals. Therefore, we favor the second alternative: that *optix* is necessary for bristle suppression by *Ubx* in the haltere.

### Forced expression of *optix* throughout the wing disc results in extravenation in the anterior wing, but does not affect the rest of the organ

The fact that *optix* expression was restricted to the anterior-most region of the wing disc made us ask whether *optix* might affect wing development if ectopically expressed throughout the developing wing. We tested this by driving a UAS-optix transgene using *nub-GAL4.* Wings of *nub>optix* adults showed extravenation in the margin cell – precisely the region where *optix* is normally expressed and required (Figure 7). However, the rest of the wing remained unaltered. This lack of effect was particularly unexpected: Optix is a transcription factor and we would have predicted that, similarly to what happens in the anterior-most wing, cells elsewhere in the pouch would have responded to its ectopic expression. This to us suggests that most parts of the wing pouch are “protected” from the action of optix, either by lack of available DNA target sequences (an epigenetic effect) or the absence of a positive co-factor (or the presence of a repressor).

**Figure 7.**
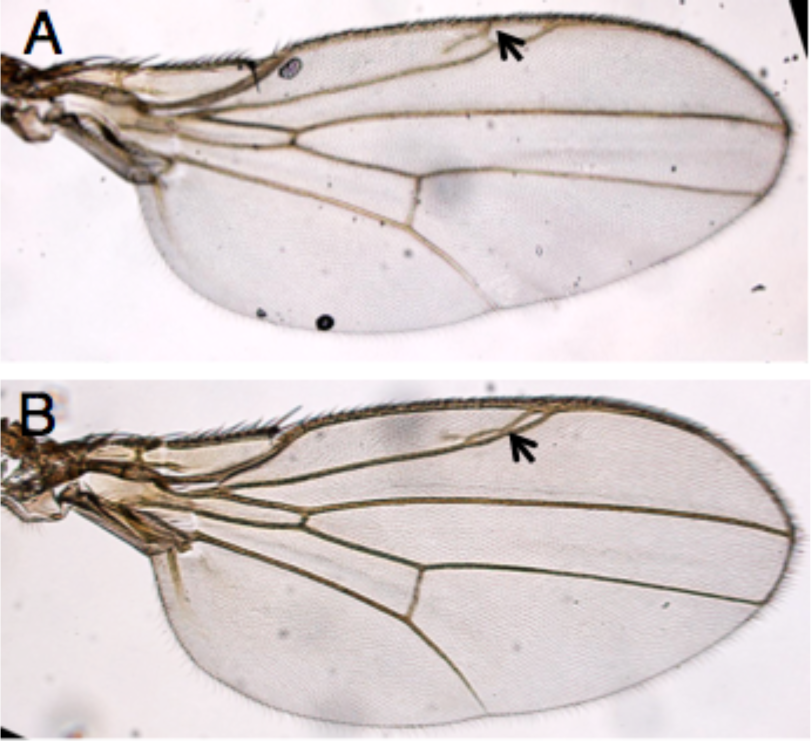
Overexpression of *optix* throughout the wing pouch causes extravenation between L1 and L2. Wings from adult male (A) and female (B) *nub>optix* flies. Arrows point to extravenation.

## DISCUSSION

The Dpp signaling gradient is required for the patterning of the whole wing, from the center to its margin (Lecuit et al., 1996; Nellen et al., 1996; Posakony et al., 1990). This gradient is translated into a series of contiguous domains expressing distinct transcription factors, each required for the specification of specific features in the adult organ (Lawrence and Struhl, 1996; Restrepo et al., 2014). However, while the transcription factors acting in the central wing were known, the most anterior region of the wing –the region comprised between the longitudinal vein 2 (L2) and the anterior margin (L1)-lacked a specific transcription factor. In this paper, we show that this transcription factor, or at least one of them, is *optix.*

Our results indicate that *optix* is expressed in, and required for the growth of this most anterior sector of the wing, the so-called margin cell. This role is in agreement with previous results showing that Six3 regulates cell proliferation in vertebrate systems (Del Bene et al., 2004; Gestri et al., 2005). We further show that Dpp signaling plays a major role in setting the *optix* expression domain. Although it has been reported before that *sal* genes are required to set the central limit of this domain, in discs lacking *sal* function *optix* does not extend all the way to the AP border (Organista et al., 2015), suggesting additional mechanisms involved in *optix* repression. The fact that *sal* is not expressed in the haltere pouch and still *optix* does not extend all the way to the AP border, the exclusion of *optix* expression from intermediate/high Dpp signaling in both wing and haltere, and the requirement of Dpp signaling to repress *optix* in any position of the anterior wing compartment globally suggested to us that either Dpp activates a different repressor closer to the AP border, or that Dpp signaling represses directly *optix* transcription. Our work cannot distinguish between these possibilities. Regarding another well characterized Dpp target, *omb*, the extensive coexpression of *omb* and *optix* in the haltere also seems to exclude *omb* as a repressor. Therefore, either another unknown repressor exists, or Dpp signaling acts as a direct *optix* repressor. While in the haltere, the domain of *optix* would be set directly by Dpp, in the wing *sal* would be an additional repressor. By intercalating *sal*, the Dpp positioning system may be able to push the limit of *optix* expression farther away from the AP border of the wing. The Sal proteins have been previously shown to act as transcriptional repressors of *knirps (kni)* to position vein L2 (de Celis and Barrio, 2000). Thus, adding *sal* repression may help to align the *optix* domain with L2. This additional repression would not be operating in the haltere, which lacks venation.

Interestingly, the logic of *optix* regulation by Dpp is different from that of other Dpp targets. The activation of the *sal* paralogs *(sal-m* and *sal-r)* and *aristaless (al)*, another target required for vein L2 formation (Campbell and Tomlinson, 1998), proceeds through a double repression mechanism: In the absence of signal, the Brinker repressor keeps *sal* and *al* off. Activation of the pathway leads to the phosphorylation of the nuclear transducer Mad (pMad) which, in turn, represses *brk*, thus relieving the repression on *sal* and *al* (Campbell and Tomlinson, 1999; Moser and Campbell, 2005). Therefore, *optix* regulation by Dpp signaling could be more direct similarly to that of *brk.*

One interesting aspect of *optix* function is that it plays qualitatively different roles in the wing and the haltere. While in the wing *optix* is required for the development of the anterior-most portion of the wing (including the margin bristles), in the haltere *optix* serves to suppress the development of sensory bristles, a task known to be carried out by the Hox gene *Ubx.* We have ruled out a role for *optix* in regulating *Ubx* expression, at least when judged from Ubx protein levels (Figure S2). Therefore, *optix* is required for a subset of Ubx’s normal functions. Since *optix* encodes a Six3-type transcription factor, this interaction could be happening at the level of target enhancers, where the combination of Ubx and Optix would allow the activation or repression of specific sets of genes.

Finally, we have observed that the expression of *optix* in wing and haltere primordia is conserved across higher Diptera (Figure 2). Interestingly, *optix* is expressed in the developing wings of passion vine butterflies (genus *Heliconius).* In *Heliconius* species, *optix* has been co-opted for red color patterning in wings (Reed et al., 2011). However, the ancestral pattern found in basal Heliconiini is in the proximal complex, a region that runs along the base of the forewing costa, the most anterior region of the forewing (Martin et al., 2014). This similarity between *optix* expression patterns in forewings of Diptera and Lepidoptera make us hypothesize that an *optix* ancestral role might have been “structural”, being required for the development of the anterior wing. Once expressed in the wings, recruitment of red pigmentation genes allowed *optix* co-option for color pattern diversification through regulatory evolution (Martin et al., 2014). We note that a pre-requisite for this co-option in wing pigmentation patterning must have been that *optix* would not interfere with the developmental pathway leading to the formation of a normal wing in the first place. The fact that the effects of *optix* overexpression throughout the wing primordium in *Drosophila* are restricted to the anterior/proximal wing -its normal expression domain-indicates that *optix* cannot engage in promiscuous gene regulation, and that its function depends on other competence factors, which would limit its gene expression regulatory potential.

## ACKNOWLEDGEMENTS

We thank P. Tomancak and C. Dahman for generously sharing RNA expression data with us, the Casares and Posnien lab members for discussions and A. lannini for technical assistance. Confocal imaging was supported by the CABD Advanced Light Microscopy facility.

## SUPPLEMENTARY FIGURES

**Figure S1.**
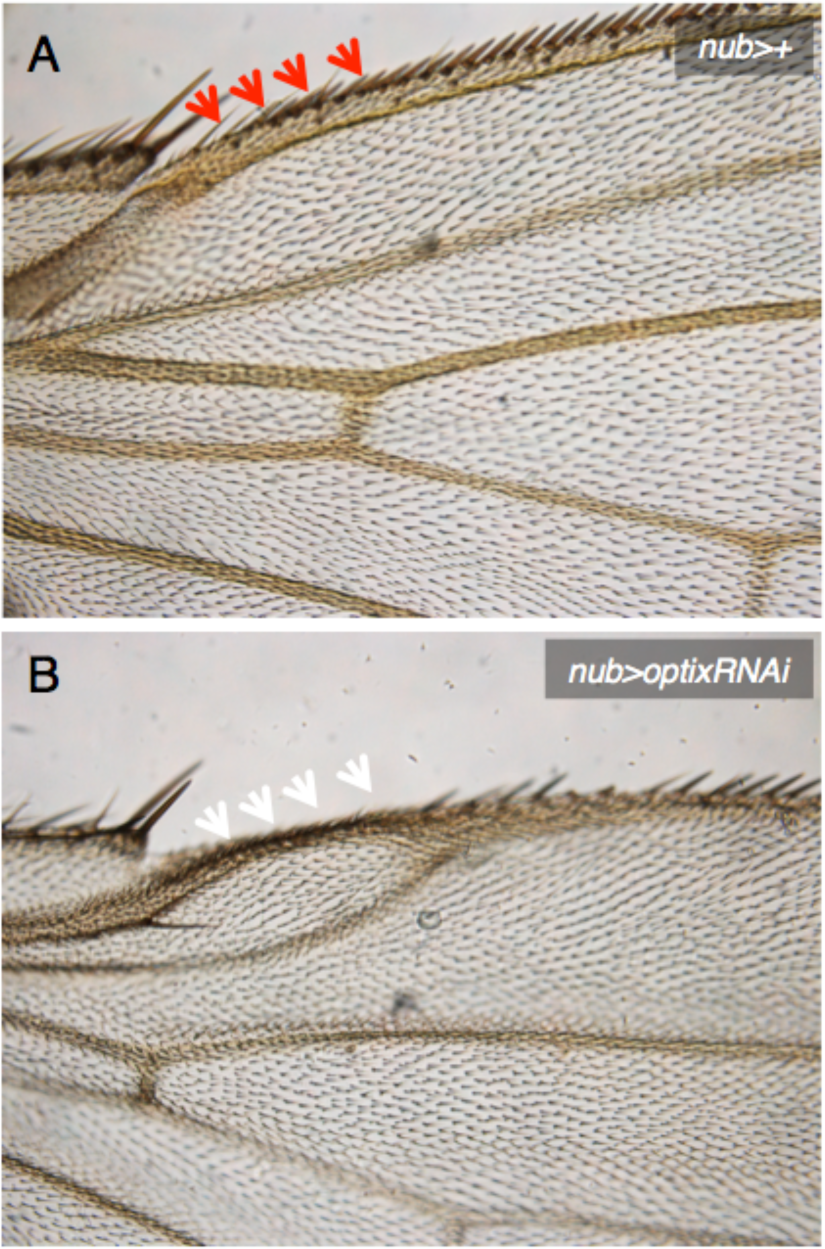
Attenuation of *optix* causes the loss of triple row sensory bristles. Control (A: *nub>)* and *optix* knock-down (B: *nub>optixRNAi).* Arrows in (A) and (B) mark equivalent positions. Note absence of bristles along the anterior-proximal margin that spans the reduced marginal cell.

**Figure S2.**
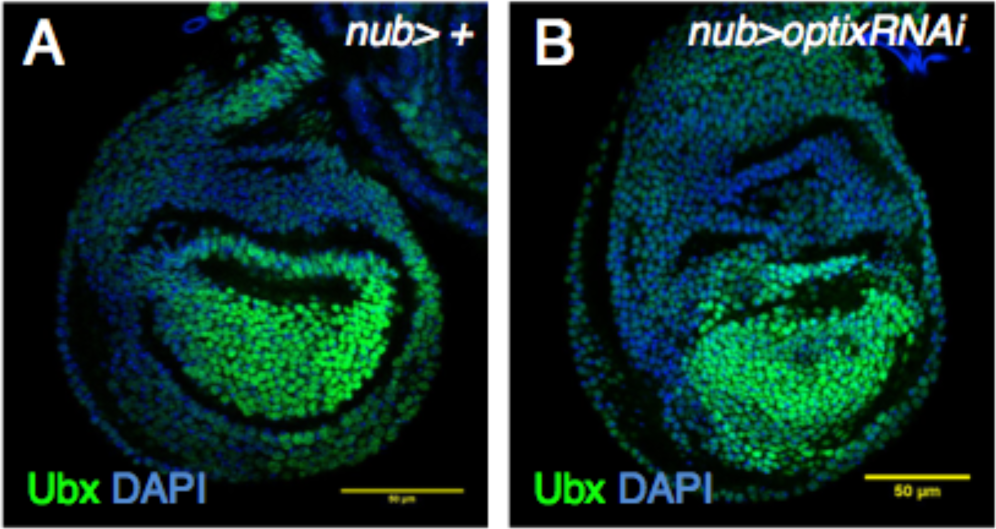
Down regulation of *optix* does not affect Ubx expression in the haltere pouch. Ubx expression in (A) control *(nub>+)* and (B) *nub>optixRNAi* haltere discs. Strong Ubx signal is detected in the haltere pouch in both situations. The discs have been counterstained with the nuclear marker DAPI.

